# Subacute Effects of Ketamine on Neural Correlates of Reward Processing

**DOI:** 10.1101/2025.09.10.675318

**Authors:** E Briem, P Stöhrmann, G Dörl, C Milz, G Schlosser, M Klöbl, M Reed, M Atger, C Schmidt, M Kathofer, GM Godbersen, JS Crone, R Lanzenberger, M Spies

**Affiliations:** Department of Psychiatry and Psychotherapy, Medical University of Vienna, Vienna, Austria; Comprehensive Center for Clinical Neurosciences and Mental Health (C3NMH), Medical University of Vienna, Vienna, Austria; Vienna Cognitive Science Hub, University of Vienna, Vienna, Austria; Faculty of Psychology, University of Vienna, Vienna, Austria

**Author notes:** Correspondance to: Prof. Rupert Lanzenberger, MD PhD, Department of Psychiatry and Psychotherapy, Medical University of Vienna Waehringer Guertel 18-20, Vienna, 1090, Austria. For submission to bioRxiv.

**Keywords:** ketamine, norketamine, fMRI, MID task, reward processing

## Abstract

**Objectives:** Ketamine’s prohedonic properties have been linked to enhanced reward-related brain activation during the early post-infusion phase. Its effects during the subacute period (∼2-24 h post-infusion), when psychotomimetic symptoms fade and neuroplastic adaptations emerge, are less well characterised. This study assessed ketamine’s subacute effects on reward processing using the Monetary Incentive Delay (MID) task.

**Methods:** In a randomised, placebo-controlled, crossover study, 28 healthy participants received 0.5 mg/kg racemic ketamine or placebo via 40-minute intravenous infusion. Functional magnetic resonance imaging (fMRI) was acquired ∼5 h post-infusion. Plasma concentrations of ketamine and norketamine were obtained for individual area under the curve (AUC) estimation. Analyses focused on the contrast between expected and actual trial outcomes.

**Results:** At five hours post-infusion, ketamine did not significantly modulate MID task-related brain activation, despite pronounced subjective drug effects. Pharmacokinetic modelling confirmed expected ketamine and norketamine profiles, but neither drug exposure (AUC) nor subjective measures correlated with neural activation.

**Conclusions:** Prohedonic effects of ketamine may not sufficiently manifest in MID task-related activation in healthy individuals ∼5 hours after infusion. The lack of significant effects provides valuable extension of the existing literature, as ketamine’s effects might be confined to a more acute time window or differ in clinical populations.

## Introduction

Ketamine’s rapid-acting antidepressant effects have led to a major shift in psychiatric research and the treatment of Major Depressive Disorder (MDD) in recent years. Unlike traditional monoaminergic antidepressants, ketamine elicits rapid antidepressant effects within 2 h and sustained effects lasting up to 7 days following a single subanaesthetic dose (Zarate et al. 2006; Coyle and Laws 2015). One of ketamine’s most promising properties is its potential to alleviate anhedonia (Lally et al. 2014; Nogo et al. 2022; Kwaśny, Kwaśna, et al. 2024; Kwaśny, Cubała, et al. 2024), a core symptom of MDD characterised by the reduced ability to experience pleasure (Gorwood 2008; Höflich et al. 2018). Anhedonia is of particular clinical interest as it correlates with poorer treatment outcomes (Spijker et al. 2001) and heightened risk of suicidality (Winer et al. 2014). Prior findings suggest that ketamine’s prohedonic effects are associated with increased glucose metabolism in reward-related brain regions, specifically the ventral striatum (Nugent et al. 2014; Lally et al. 2015).

At a neurobiological and network level, anhedonia is hypothesised to reflect deficits in reward circuitry (Gorwood 2008; Argyropoulos and Nutt 2013; Höflich et al. 2018; Pulcu et al. 2022). Reward processing relies on coordinated activity across the striatum, insula, amygdala, thalamus, and dopaminergic mesocorticolimbic regions (Oldham et al. 2018), enabling adaptive decision-making and goal-directed behaviour (Balleine and O’Doherty 2010). The Monetary Incentive Delay (MID) task is a well-established paradigm for investigating reward processing, predominantly combined with functional magnetic resonance imaging (fMRI) (Knutson et al. 2000; Lutz and Widmer 2014; Oldham et al. 2018; Wilson et al. 2018) and has been previously applied in our laboratory (Hahn et al. 2021; Hahn et al. 2025). The MID task consists of two phases: anticipation, when a cue signals potential reward or loss, and outcome, when the reward or loss is obtained or omitted. Anticipation activates regions such as the striatum, insula, thalamus, and amygdala (Knutson et al. 2000; Oldham et al. 2018; Wilson et al. 2018). Similar regions are also active during the outcome phase, suggesting overlapping neural circuits in different stages of reward processing (Knutson et al. 2000; Oldham et al. 2018; Wilson et al. 2018).

Ketamine, originally developed as an anaesthetic (Domino et al. 1965), elicits rapid antidepressant effects after a 40-minute subanaesthetic infusion (0.5 mg/kg), with improvement of depressive symptoms emerging within hours (Berman et al. 2000; Zarate et al. 2006; Nikolin et al. 2023). While its anaesthetic and psychotomimetic properties are mediated by NMDA receptor antagonism, antidepressant actions likely involve downstream mechanisms such as AMPA receptor activation and synaptic plasticity (Zanos et al. 2016; Zanos et al. 2018; Lumsden et al. 2019; Casarotto et al. 2021). Ketamine’s pharmacokinetic profile is characterised by rapid distribution, extensive hepatic metabolism, and an elimination half-life of 2 to 4 h (Mion and Villevieille 2013; Peltoniemi et al. 2016). Norketamine, the primary active metabolite, peaks around 100 min post-infusion (Zanos et al. 2016) and also exhibits NMDA receptor antagonism, although with lower affinity than ketamine itself (Ebert et al. 1997; Sałat et al. 2015).

While acute modulatory effects of ketamine on reward processing have been observed 40 min post-infusion in healthy subjects (Francois et al. 2016) and 2 h post-infusion in remitted MDD patients (Kotoula et al. 2022), subacute effects remain underexplored. In this study, we aim to assess the subacute effects of ketamine (∼5 h post-infusion) on reward circuitry using task-based fMRI during the MID task. Investigating the intermediate phase may provide unique insights into ketamine’s neuromodulatory effects following the resolution of acute psychotomimetic symptoms and during the early stages of synaptic and circuit-level reorganization. Studying healthy participants allows investigation of ketamine’s effects on reward processing without the confounds of clinical symptoms. Based on previous fMRI findings (Francois et al. 2016; Mkrtchian et al. 2021; Kotoula et al. 2022), we hypothesise that ketamine will increase neural activation in the striatum during reward anticipation, potentially reflecting its prohedonic properties (Lally et al. 2014; Nugent et al. 2014; Lally et al. 2015; Mkrtchian et al. 2021; Nogo et al. 2022; Pulcu et al. 2022).

## Methods

### Participants

Participants were required to be 18-55 years old, right-handed, in good general and psychiatric health (based on medical history, physical exam, and structured DSM-5 interview), and able to provide informed consent. Participants were excluded in case of current or past psychiatric diagnosis, relevant medical illness, pregnancy or breastfeeding, MRI contraindications, substance abuse, or prior ketamine use. Urine drug screening and pregnancy testing (in females) were performed at each visit.

### Study Design

This study was approved by the ethics committee of the Medical University of Vienna, Austria (EK 1572/2021) and performed in accordance with the declaration of Helsinki.

The study followed a single-blind, placebo-controlled, crossover design consisting of four sessions: a screening visit, two experimental sessions (ketamine and placebo in a randomised order, both including MRI scanning; Figure 1), and an end-of-study visit. At both MRI sessions, subjects received either placebo (0.9% saline solution) or an infusion of 0.5 mg/kg racemic ketamine diluted in 0.9% saline solution via a venous catheter using a continuous infusion pump over 40 min. The study medication (50 mg/mL ampoules; Hameln Pharma Plus GmbH) was prepared by a medical doctor before each session. Participants were observed in a clinical setting for at least 90 min post-infusion until full recovery from ketamine-induced sedative and dissociative effects.

**Figure 1.**
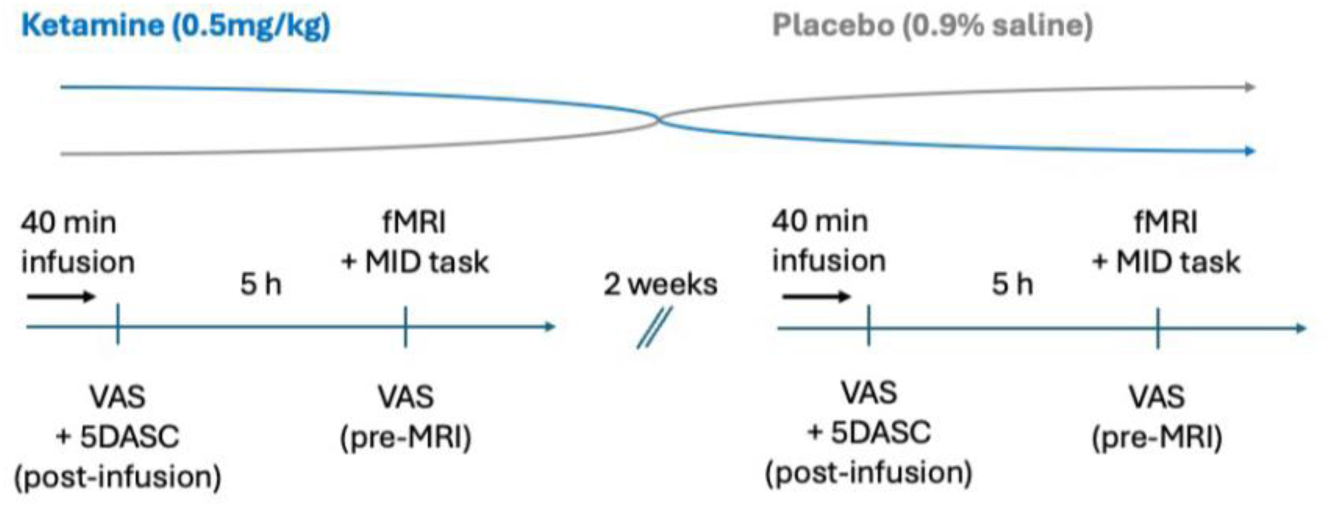
Single-blind, placebo-controlled, crossover design study design. Participants received either ketamine (0.5 mg/kg, 40-min i.v. infusion) or placebo (0.9% saline) in two separate sessions, spaced 2 weeks apart. Following each infusion, subjective drug effects were assessed using a visual analogue scale (VAS) to assess common ketamine-related adverse effects and a 5D-ASC questionnaire immediately post-infusion. Approximately 5 h later, participants underwent fMRI scanning while performing the Monetary Incentive Delay (MID) task, with VAS ratings collected immediately before scanning.

### Psychometric Measures

Subjective drug effects as experienced during ketamine application were assessed using the 5-dimensional altered states of consciousness rating scale (5D-ASC), a validated questionnaire designed to assess altered states of consciousness induced by psychoactive substances such as ketamine (Dittrich 1998; Studerus et al. 2010). Participants completed the questionnaire on a tablet in a quiet room under supervision. The 94 items of the 5D-ASC were scored on a scale ranging from 0 (no experience) to 100 (strongest possible experience). A total 5D-ASC score was computed by summing the scores of all 94 items. In addition to the 5D-ASC, participants completed two brief visual analogue scale (VAS) assessments to evaluate the duration and intensity of common ketamine-associated adverse effects, including dizziness, blurred vision, headache, nausea, dry mouth, poor coordination, and restlessness. (Murrough et al. 2013). Each of the seven symptoms was rated individually from 0 (“no symptom”) to 100 (“most severe”), and a total VAS score was obtained by summing the ratings across all symptoms. VAS assessments were conducted shortly after infusion stop as well as immediately before MRI scanning.

### fMRI Scanning

Task-based BOLD fMRI scanning was conducted approximately 5 h (mean: 5 h 19 min ± 6.5 min) after the start of the ketamine infusion. MRI scanning was performed on a 3 Tesla MR Scanner (MAGNETOM Prisma, Siemens Medical, Erlangen, Germany). Task-based fMRI data was acquired using a single-shot gradient-recalled echo-planar imaging (EPI) sequence with the following parameters: echo time (TE) / repetition time (TR) = 30 ms / 1600 ms, matrix size of 70 × 70 voxels, a field of view (FOV) of 210 × 210 mm, and a multiband factor of 2. The resulting voxel dimensions were 3 x 3 x 3 mm, with a total of 48 slices (slice thickness = 3 mm). Preprocessing included conversion of data to NIfTI-format, physiological noise correction (PESTICA; Shin et al. 2022), motion correction (SLOMOCO; Beall and Lowe 2014)), slice time correction (SPM12), wavelet despiking (Patel et al. 2014; Patel and Bullmore 2016), distortion correction (TOPUP; Andersson et al. 2003), spatial normalisation to MNI space via the mean T1 image across sessions (SPM12), and spatial smoothing using a 9 mm FWHM Gaussian kernel.

### MID Task

During the MID task, participants were instructed to maximise gains and avoid losses during win, lose, and neutral trials. Each trial began with the presentation of a cue (anticipation phase) indicating a potential gain or loss (e.g., –1€, +3€). After a variable delay of 3–5 seconds, a target stimulus appeared (“!”), prompting participants to press a button as quickly as possible. If the reaction time was within a predefined time window based on their individual average response time determined prior to the scan, the amount was gained or the loss was avoided; otherwise, the amount was not gained, or the loss was incurred. Each response was immediately followed by feedback displaying the outcome and the updated cumulative balance (outcome phase). Unknown to the participants, odds were manipulated during “win” and “loss” trials by increasing or decreasing the valid time window for reaction, respectively. The task lasted 10 min and 5 s and consisted of nine blocks presented in randomised order (three gain, three loss, three neutral e.g., 0 € gambled), comprising a total of 57 trials (3 × 7 gain, 3 × 7 loss, 3 × 5 neutral). Trials were separated by a fixation cross presented for a variable interval of 3– 7 seconds. To maintain attention and enable modelling of reward sensitivity, six different monetary amounts were used (0.5€, 1€, and 3€ for both gain and loss trials; initial balance: 10€). Monetary amounts were selected to reflect a range of low to high incentives, in line with previous studies on motivational salience (Knutson et al. 2000; Kotoula et al. 2022). The final balance (if positive) was paid out to participants at the end of the study.

### Pharmacokinetic Analysis

Racemic ketamine (0.5 mg/kg) was administered intravenously over 40 min, and blood samples were collected at predefined time points (0, 5, 10, 20, 40, 100, and 360 min). After centrifugation and plasma separation, plasma samples were stored at ≤−20°C until analysis. Ketamine plasma concentrations were determined by gas chromatography– tandem mass spectrometry with multiple reaction monitoring at the Department of Laboratory Medicine, Medical University of Vienna, Austria. The analytical procedure was validated in accordance with EMA guidelines (Bioanalytical method validation - Scientific guideline | European Medicines Agency (EMA) 2011 Aug 1).

Area under the curve (AUC) values for ketamine and its primary metabolite, norketamine, were calculated from model-fitted concentration-time profiles. MATLAB’s SimBiology toolbox was used to estimate individual concentration trajectories based on subject-specific pharmacokinetic (PK) parameters, enabling quantification of systemic drug exposure by accounting for inter-individual variability in drug distribution and clearance. The PK analysis was performed using a two-compartment model (central and peripheral) with two metabolic delay compartments to simulate the conversion of ketamine to norketamine. The model structure was based on the PK framework described by Kamp et al. (Kamp et al. 2020), which applied a compartmental approach to account for the distribution and metabolism of ketamine and its metabolites. Parameter estimation was carried out using nonlinear mixed-effects modelling (NLME) in SimBiology (MATLAB). Individual pharmacokinetic parameters such as clearance (CL), volume of distribution (V_central_, V_peripheral_), and inter-compartmental transfer rates (Q) were estimated from the model based on the observed concentration-time data. Model validation involved visual comparison of simulated and observed plasma concentrations of both ketamine and norketamine.

### fMRI Analysis and Statistical Modelling

#### First Level Analysis

At the first level, each participant’s data was modelled using a general linear model (GLM). Pretrial cues (win/loss/neutral), targets, and feedback (gain/no gain/loss/no loss/neutral) were modelled separately - with and without parametric modulation by reward magnitude (absolute value, log- or rank-transformed to test for different neural response profiles). To control for residual physiological and other sources of noise, principal components of the white matter and CSF signal time courses were included as nuisance regressors (CompCor; Behzadi et al. 2007).

In order to capture neural processes associated with the alignment or mismatch between anticipated and actual outcomes (Knutson and Greer 2008; Schultz 2016), we additionally implemented an Expectation–Outcome Congruency contrast as a measure of reward prediction error. This contrast compared trials in which the actual feedback matched the expected outcome (congruent: potential win/loss was signalized and coincided with feedback “win” or “loss”) versus trials where the outcome violated expectations (incongruent: no gains/no losses), thereby extending the traditional MID analysis beyond separate anticipation and outcome. We speculated that psychopathologies (e.g. MDD), their symptoms (anhedonia), and substances affecting reward circuitry (ketamine) might influence not only specific aspects of win and loss processing (e.g. the time window in which positive or negative feedback is received) but also the evaluation of outcomes relative to expectations — that is, how “best” or “worst” anticipated outcomes are perceived compared with the actual experience.

#### Second Level Analysis

##### Region of Interest (ROI)-based Analysis

We conducted a literature-driven, a priori ROI-based analysis, focusing on key brain structures implicated in reward and salience processing, including the striatum, nucleus accumbens (NAc), insula, amygdala, and thalamus (Knutson and Greer 2008; Dugré et al. 2018; Oldham et al. 2018; Jauhar et al. 2021; Chen et al. 2022). ROIs were anatomically defined using the Automated Anatomical Labeling 3 (AAL3) atlas (Rolls et al. 2020). For each ROI, contrast estimates (beta values of the first level analysis) were extracted and analysed using linear mixed-effects models in MATLAB to examine the influence of ketamine exposure, subjective drug experience, and covariates on ROI activation.

The primary model specification was:

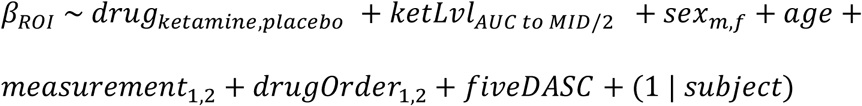

Here, ketLvl represents the area under a piecewise linear approximation of individual ketamine exposure up to half the task duration (z-scored within the ketamine condition, otherwise zero, to avoid collinearity with the drug conditions). fiveDASC reflects subjective drug experience (5D-ASC, z-scored within each condition). A random intercept for subject was included to account for repeated measures.

##### Whole-Brain Analysis

In addition to the ROI-based approach, on an exploratory basis, the same first-level contrasts were entered into second-level whole-brain analyses using a flexible factorial design in SPM12. The model accounted for drug (placebo/ketamine), measurement, order of drugs (to account for potential carry-over and/or habituation effects), and enabled the testing of main and interaction effects using F- and t-tests. Findings were considered significant if p_uncorr_ < 0.001 and p_FWE, cluster_ < 0.05.

## Results

A total of 36 healthy subjects completed the study. After excluding subjects with either excessive head motion during MRI scanning, missing plasma concentration values or irregular ketamine plasma concentration curves (e.g. peak plasma concentration 20 minutes prior to end of infusion), 28 participants (15 females, mean age 23.6 ± 3.8 years) remained eligible for analysis.

### Pharmakokinetic Data

Ketamine plasma levels increased rapidly after infusion, peaking at approximately 40 min, i.e., end of infusion (Figure 2A, Table 1,), while norketamine concentrations increased more slowly, consistent with the expected metabolic conversion over time (Figure 2B, Table 2). AUC values for ketamine and norketamine plasma concentration curves were calculated up to half the task duration (5 h 14 min ± 6 min) using a two-compartment PK model (SimBiology, MATLAB) and model-fitted concentration-time profiles (Figure 3).

**Figure 2.**
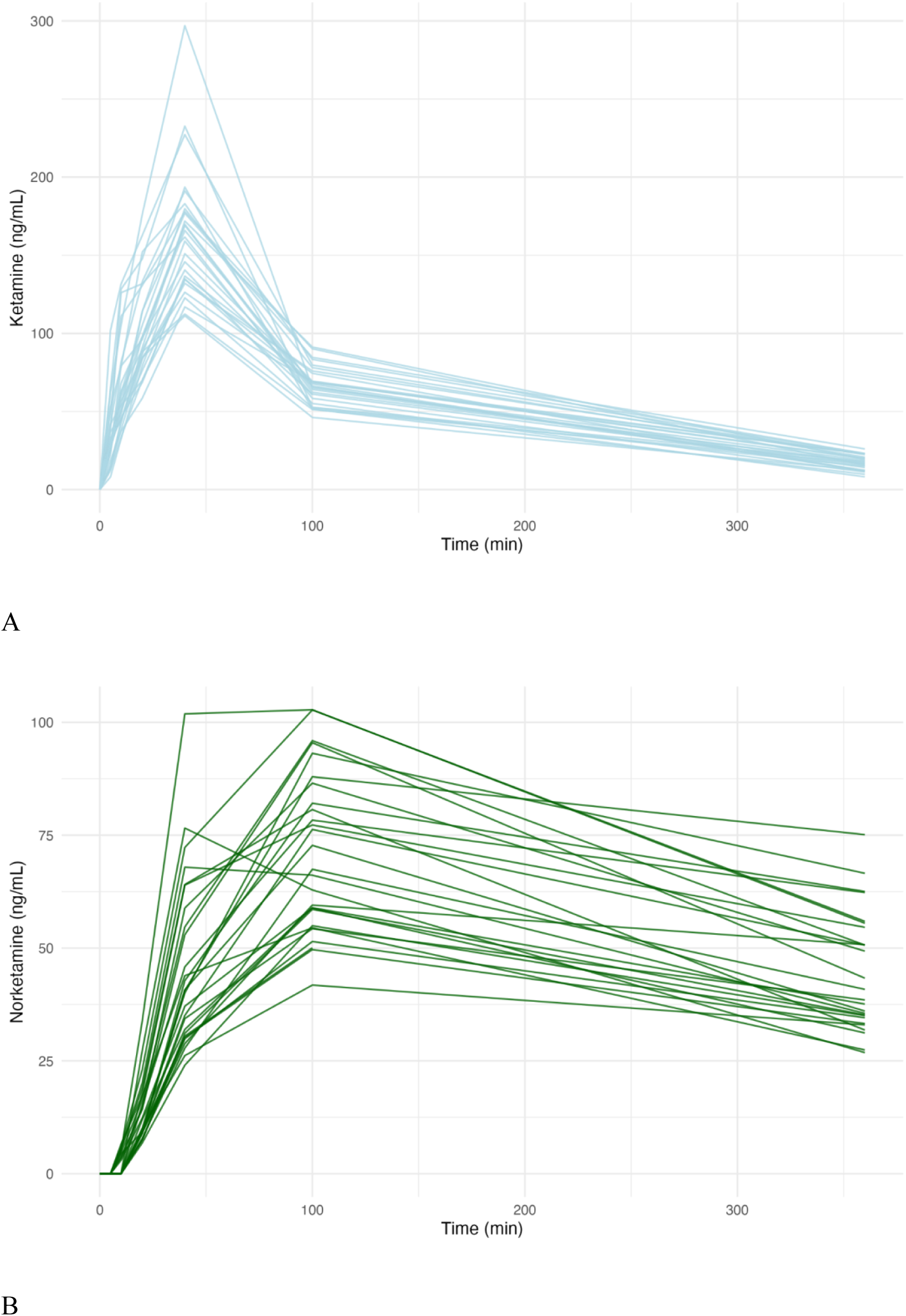
Plasma concentration–time curves of ketamine (A, blue) and norketamine (B, green) following a 40-min intravenous infusion of racemic ketamine (0.5 mg/kg). Each line represents an individual subject (n = 28). Ketamine concentrations peak at infusion stop, while norketamine shows a delayed peak due to metabolic conversion.

**Figure 3.**
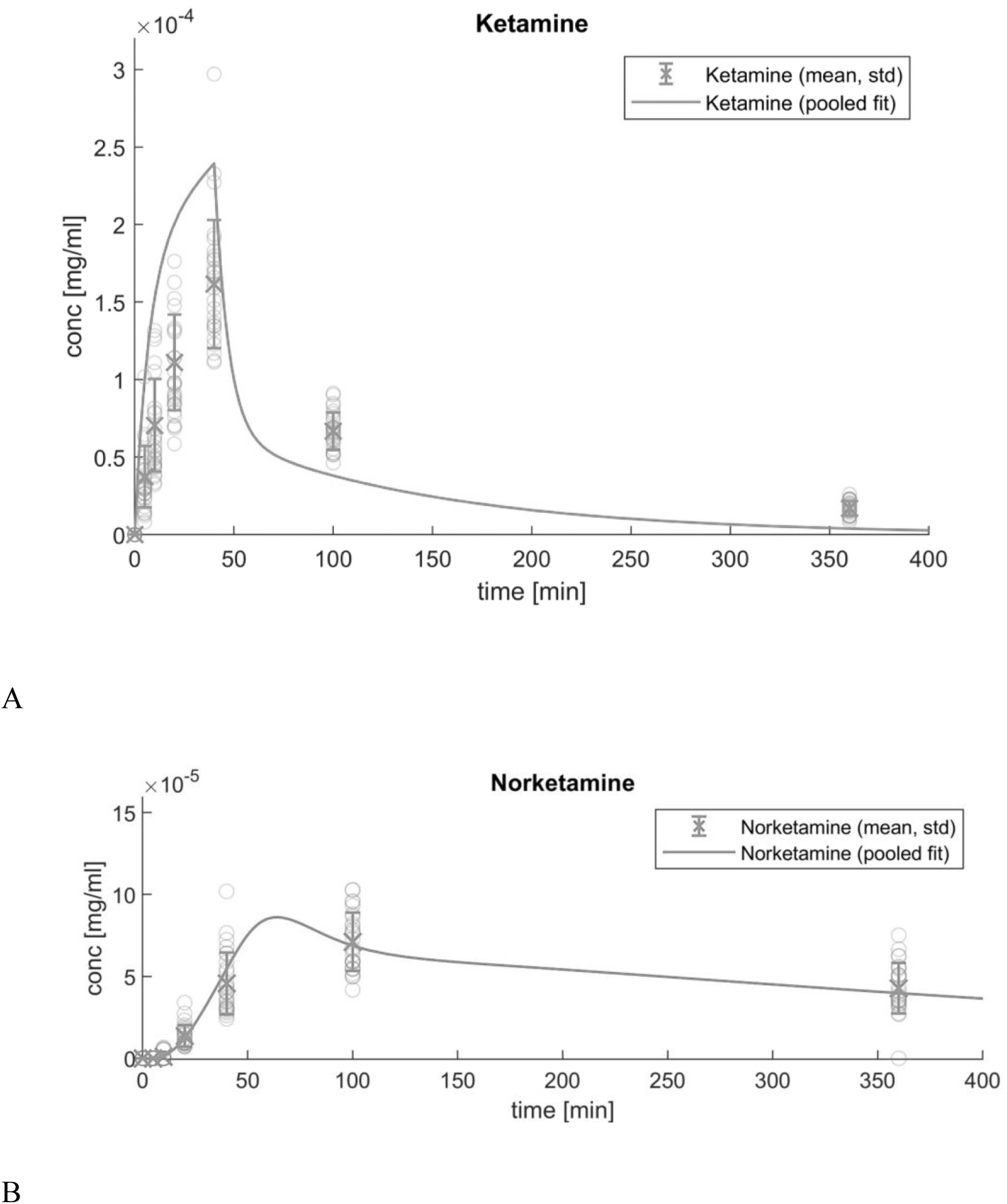
Piece-wise, linearly fitted plasma concentration–time curves for ketamine (A) and norketamine (B) following a 40-min intravenous infusion of racemic ketamine (0.5 mg/kg). Symbols (open circles) represent individual data points and mean ± SD (crosses with error bars), while solid lines depict the pooled pharmacokinetic model fit.

**Table 1.** Plasma concentrations of ketamine (n = 28). Mean, standard deviation, minimum, and maximum plasma levels of ketamine at each time point following a 40-minute intravenous ketamine infusion (0.5 mg/kg, infusion start = 0min).

| Time (min) | Ketamine_Mean (ng/ml) | Ketamine_SD (ng/ml) | Ketamine_Min (ng/ml) | Ketamine_Max (ng/ml) |
| --- | --- | --- | --- | --- |
| 0 | 0.0 | 0.0 | 0.0 | 0.0 |
| 5 | 39.08 | 31.01 | 8.08 | 166.7 |
| 10 | 73.05 | 40.62 | 32.43 | 186.44 |
| 20 | 115.01 | 52.48 | 58.55 | 285.85 |
| 40 | 165.19 | 41.81 | 111.26 | 297.0 |
| 100 | 67.18 | 12.09 | 46.21 | 91.27 |
| 360 | 17.19 | 4.61 | 8.2 | 26.05 |

**Table 2.** Plasma concentrations of norketamine (n = 28). Mean, standard deviation, minimum, and maximum plasma levels of norketamine at each time point following a 40-minute intravenous ketamine infusion (0.5 mg/kg, infusion start = 0min).

| Time (min) | Norketamine_Mean (ng/ml) | Norketamine_SD (ng/ml) | Norketamine_Min (ng/ml) | Norketamine_Max (ng/ml) |
| --- | --- | --- | --- | --- |
| 0 | 0.0 | 0.0 | 0.0 | 0.0 |
| 5 | 0.0 | 0.0 | 0.0 | 0.0 |
| 10 | 1.93 | 2.49 | 0.0 | 6.58 |
| 20 | 15.4 | 9.89 | 6.84 | 49.35 |
| 40 | 45.63 | 18.68 | 24.04 | 101.88 |
| 100 | 70.85 | 17.43 | 41.79 | 102.78 |
| 360 | 44.4 | 13.06 | 26.83 | 75.12 |

### Subjective Effects

5D-ASC and VAS scores were available for 27 of the 28 subjects. Total 5D-ASC scores were significantly elevated during ketamine visits (2969.8 ± 1363.3) compared to placebo visits (251.6 ± 406.1; paired t(26) = 10.09, p < .0001, Cohen’s d = 2.67; Figure 4), indicating pronounced subjective drug effects during ketamine sessions. VAS scores immediately post-infusion were also significantly higher (281.5 ± 72.2) compared to placebo (32.8 ± 51.5; paired t(26) = 16.49, p < .0001, Cohen’s d = 3.93). During ketamine sessions, VAS scores decreased substantially from immediately post-infusion (281.5 ± 72.2) to just before MRI scanning (35.3 ± 43; paired t(26) = 18.5, p < .0001, Cohen’s d = 4). Nevertheless, compared to placebo sessions (9.3 ± 19.1), VAS scores just before MRI remained significantly higher during ketamine sessions (median difference = 26; Wilcoxon signed-rank test: V = 189, p = 0.0018; Figure 5). Distributional checks confirmed use of paired t-tests for 5D-ASC and VAS post-infusion, while non-normal pre-MRI VAS differences were analysed with the Wilcoxon signed-rank test.

**Figure 4.**
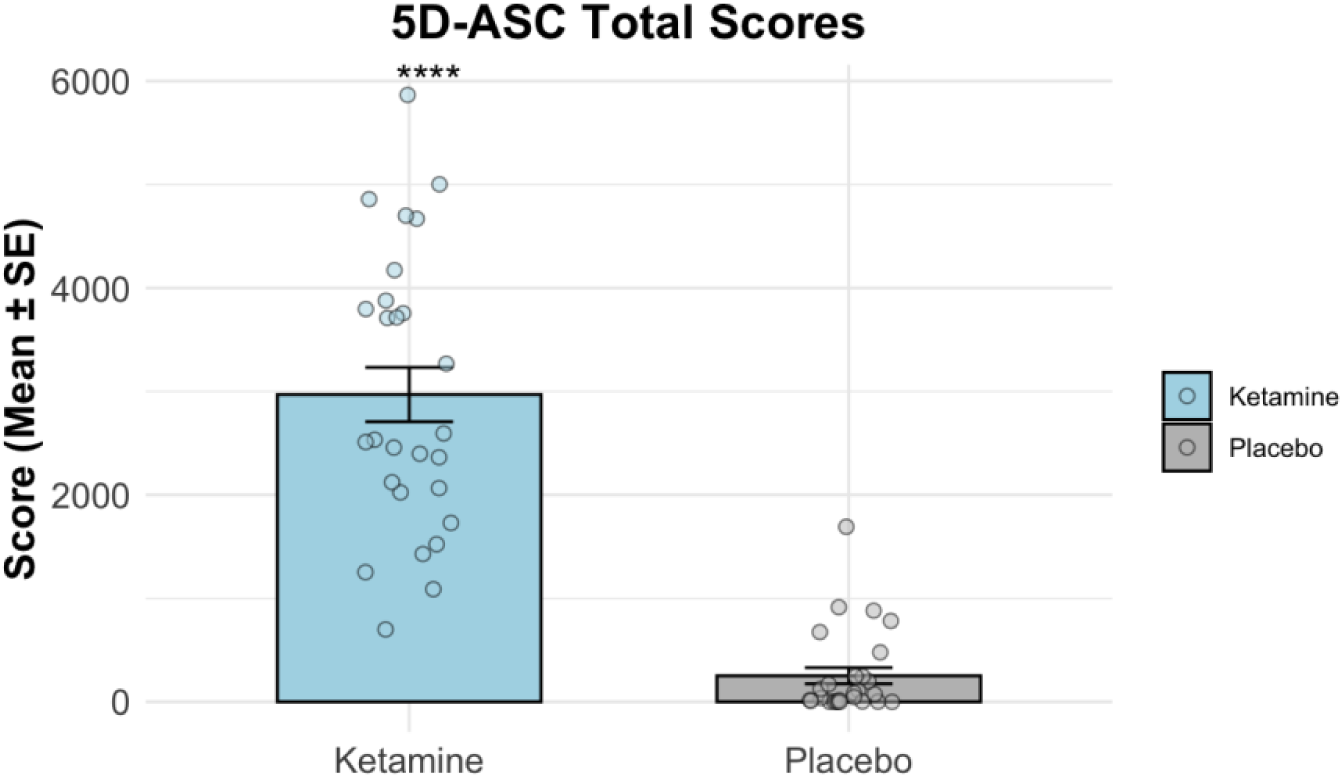
Subjective drug effects measured via the 5-dimensional altered states of consciousness rating scale (5D-ASC) questionnaire (n = 27). Total 5D-ASC scores post-infusion for the ketamine and placebo conditions. Bars represent mean ± standard deviation. Asterisks indicate significant differences (**p < .01, ****p < .0001).

**Figure 5.**
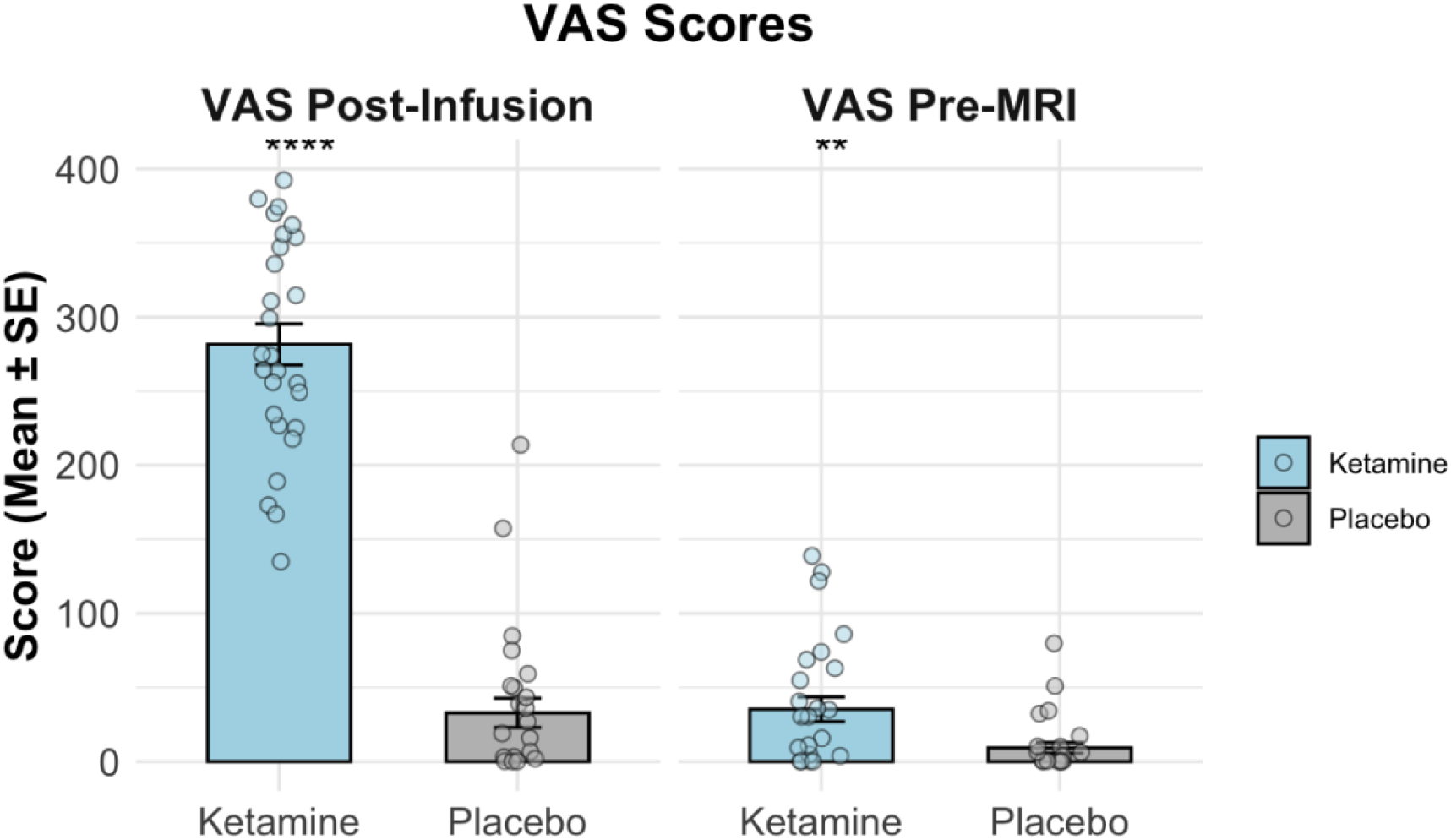
Visual analogue scale (VAS) ratings of common ketamine-associated adverse effects for the ketamine and placebo conditions (n = 27). Ratings were obtained shortly after infusion stop (post-infusion) and immediately before MRI scanning (pre-MRI). The assessed symptoms included dizziness, blurred vision, headache, nausea, dry mouth, poor coordination, and restlessness (Murrough et al. 2013). Bars represent mean ± standard deviation. Asterisks indicate significant differences between conditions (**p < .01, ****p < .0001). Group differences were tested with paired t-tests (VAS Post-Infusion) or Wilcoxon signed-rank tests (VAS Pre-MRI).

### MID Task Performance

Reaction times during the MID task did not significantly differ (α = 0.05) between ketamine and placebo condition (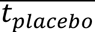≈ 338 *ms*, 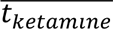≈ 352 *ms*, p = 0.07, paired t-test; 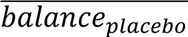 ≈ 15.82 €, 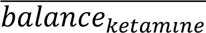≈ 15.51 €, p = 0.78, paired t-test) (Figure 6). Normality tests confirmed that assumptions for the paired t-test were met (all p > .05).

**Figure 6.**
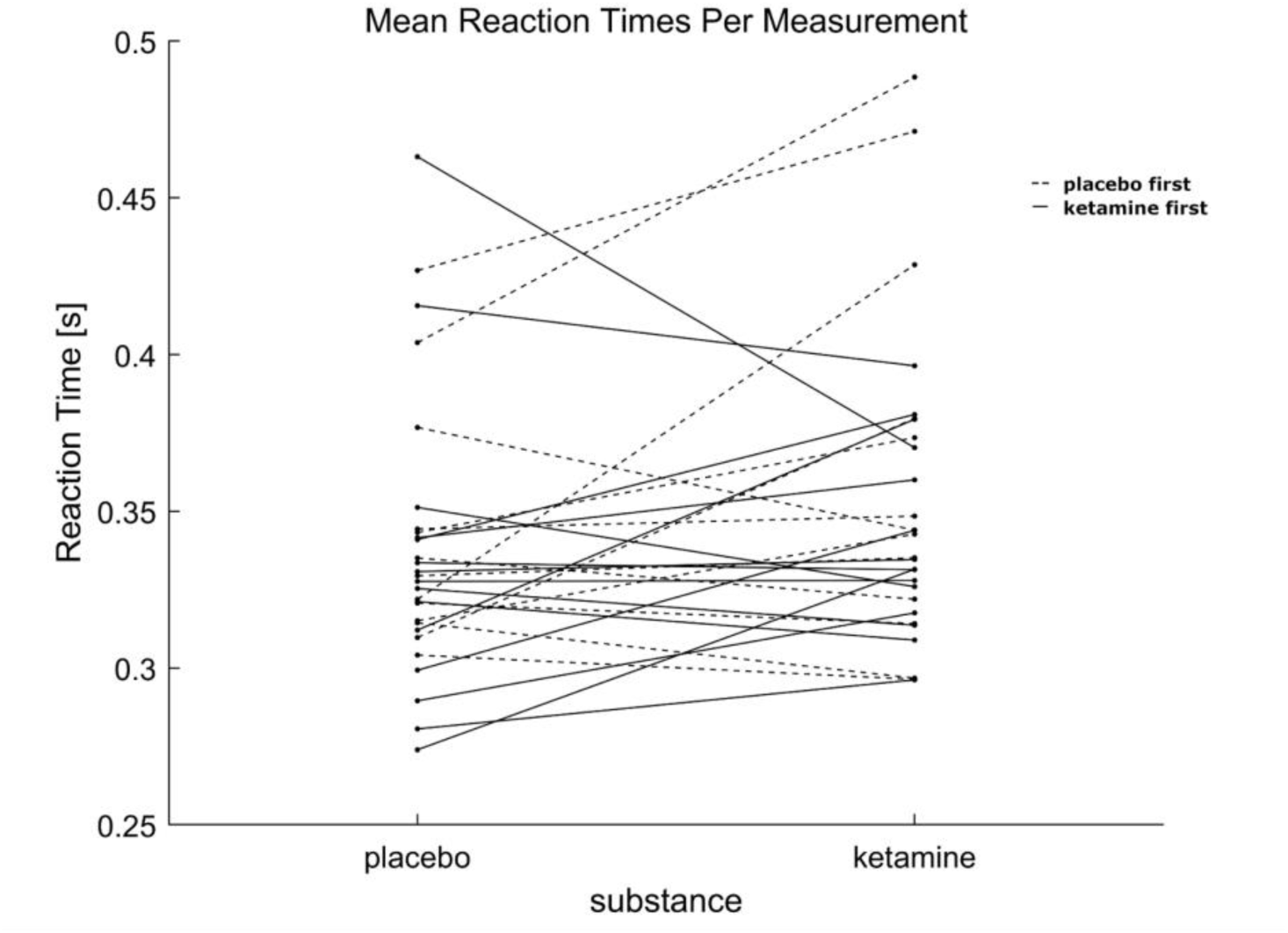
Mean reaction times during the MID task did not significantly differ between ketamine and placebo sessions, irrespective of session order.

### fMRI Analysis

#### First Level Analysis

At the single-subject level, contrasts for anticipation, feedback (win, no-win, loss, no-loss), and Expectation–Outcome Congruency elicited robust activation patterns. Representative first-level statistical maps (Figure 7) illustrate engagement of reward- and salience-related regions.

**Figure 7.**
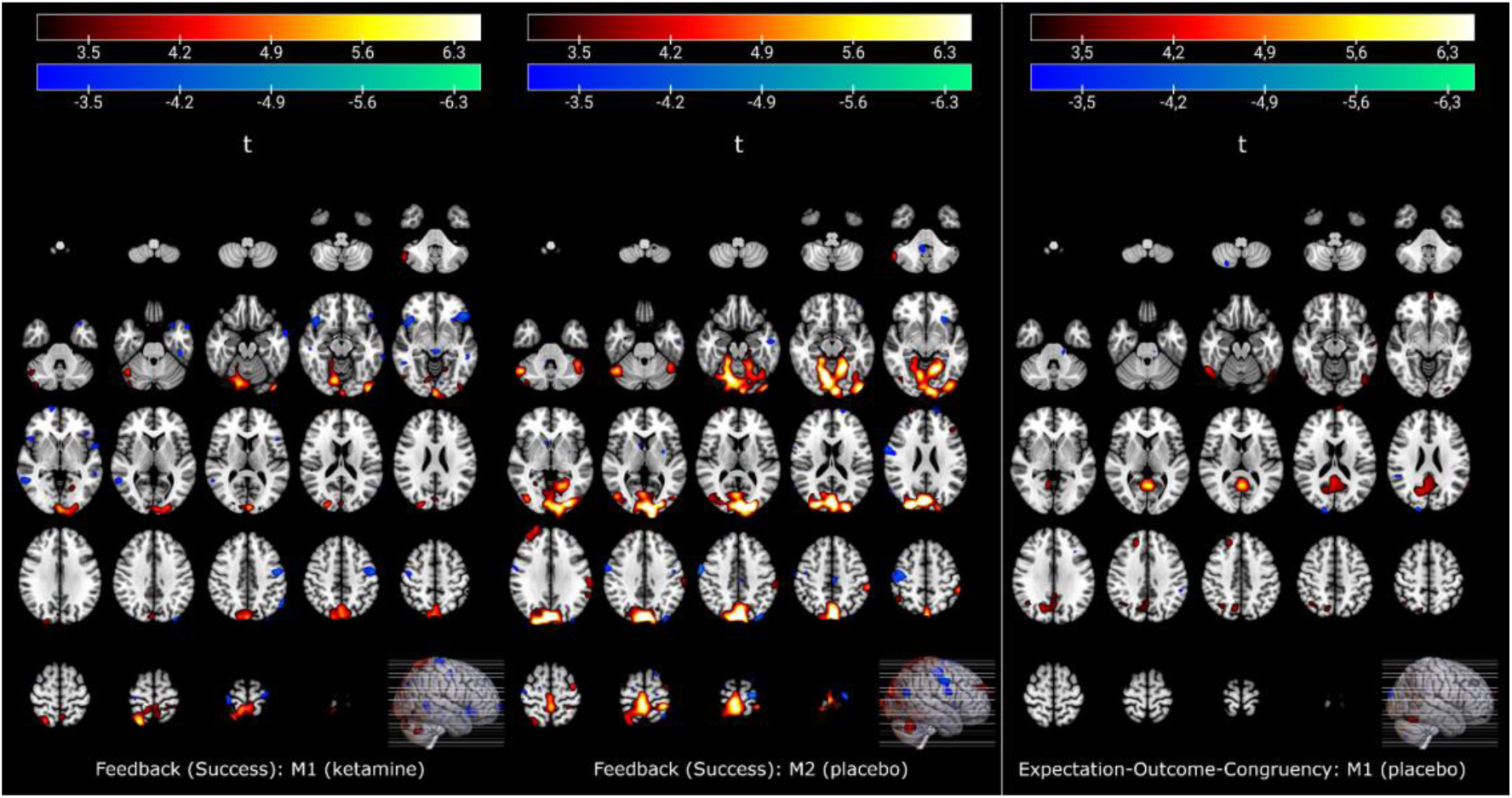
Exemplary first level whole-brain activation maps (p_uncorr_ < 0.001 shown) for feedback “success” (win or loss prevented) during the MID task under ketamine (left) and placebo (center) and expectation-outcome-congruency (right, placebo condition).

#### Second Level Analysis

##### ROI-based Analysis

ROI analyses focused on the amygdala, insula, NAc, striatum, and thalamus, based on their well-established role in reward and salience processing (Knutson and Greer 2008; Dugré et al. 2018; Oldham et al. 2018; Jauhar et al. 2021; Chen et al. 2022). For each ROI, first level contrast estimates (beta values) were extracted and analysed using linear mixed-effects modelling in MATLAB to assess the influence of drug condition (ketamine vs. placebo), individual ketamine exposure (modelled as AUC until task onset), and subjective drug experience (5D-ASC scores). No significant main effect of drug condition was observed for any ROI. In the thalamus, a significant positive association between 5D-ASC scores and BOLD activation was found (p = 0.013). However, given the absence of drug effects and multiplicity adjustments this association may represent a false positive.

##### Whole-Brain Analysis

Since the ROI-based analysis did not yield any significant results, we conducted an exploratory whole-brain analysis. Group-level statistical analysis was performed using a flexible factorial design in SPM12, which accounted for within-subject variability in the crossover design. Neither the main effect of drug condition (difference between ketamine and placebo measurement) nor exploratory F-contrasts for other factors revealed significant differences in BOLD activation. Additional exploratory models testing alternative parametrizations of monetary reward (reward magnitude, rank, or log magnitude) and feedback- or anticipation-only phases also yielded no detectable substance effects. However, exploring each substance individually, spatially coherent, significant activation patterns were found (Figure 8).

**Figure 8.**
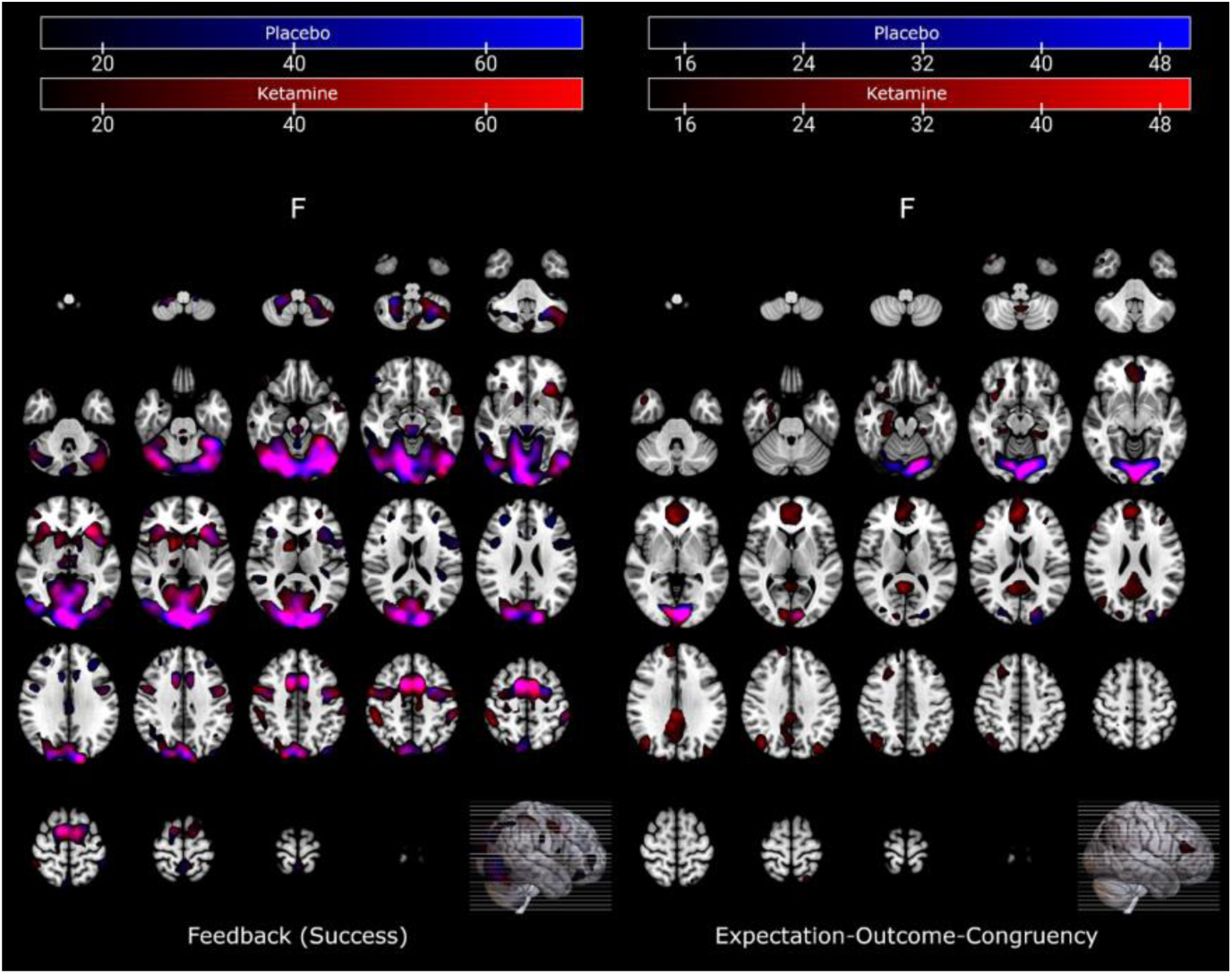
Group-level whole-brain activation maps (p_uncorr_ < 0.001 shown) for feedback “success” (win or loss prevented) during the MID task (left) and expectation-outcome-congruency (right). Activation shown was modelled without parametrization of amount gambled. Placebo condition statistics are displayed in blue, ketamine’s in red. Additive overlay blending was used for emphasis of spatial overlap between conditions.

## Discussion

Ketamine administration 5 h prior to scanning did not significantly alter reward-related brain activation during the MID task at either the ROI or whole-brain level. Pharmacokinetic modelling confirmed expected ketamine and norketamine concentration profiles, but neither individual drug exposure (AUC) nor subjective experience correlated with neural activation.

Our ROI-based analysis focused on the striatum, NAc, thalamus, insula, and amygdala. These regions were selected based on meta-analytic evidence identifying them as core neural substrates of reward and loss anticipation (Knutson and Greer 2008; Dugré et al. 2018; Oldham et al. 2018; Jauhar et al. 2021; Chen et al. 2022). While the insula, amygdala and thalamus are widely associated with motivational salience (Seeley et al. 2007; Menon and Uddin 2010; Oldham et al. 2018), the striatum, particularly the NAc, has been linked to reward magnitude (Knutson et al. 2001; Costumero et al. 2013), reward sensitivity (Carter et al. 2009; Hahn et al. 2009; Costumero et al. 2013) and the subjective experience of motivation (Knutson et al. 2001). Striatal hypoactivation, particularly within the NAc, is consistently observed in depression during reward anticipation and feedback (Pizzagalli et al. 2009; Smoski et al. 2009; Zhang et al. 2013; Admon and Pizzagalli 2015) and is considered a neural correlate of anhedonia (Szczepanik et al. 2019; Borsini et al. 2020). Given ketamine’s proposed prohedonic effects (Lally et al. 2014; Nogo et al. 2022; Pulcu et al. 2022), we hypothesised that it might modulate activity within these ROIs during the MID task. Our Expectation-Outcome Congruency contrast was designed to capture neural activity related to outcome monitoring and reward prediction errors, processes essential for adaptive learning and frequently impaired in MDD (Schultz et al. 1997; Pizzagalli et al. 2009; Smoski et al. 2009).

Our findings provide further information on the temporal effects of ketamine on reward processing when combined with previous studies. Kotoula et al. reported increased activation of the NAc, putamen, insula, and caudate in remitted MDD patients approximately 2 h post-infusion during the feedback phase of the MID task, though only in ROI analyses (Kotoula et al. 2022). François et al. reported attenuated ventral striatal response during reward anticipation in both humans and rodents within 40 min post-infusion (Francois et al. 2016). While these findings may initially appear divergent, they likely reflect differences in timing. Both studies assessed participants in the acute post-infusion phase (40min and 2h post-administration), when plasma concentrations of ketamine and norketamine were near their peak levels and NMDA receptor antagonism ongoing (Zanos et al. 2018; Kamp et al. 2020). François et al. captured the acute phase near peak NMDA receptor blockade, whereas Kotoula et al. examined a slightly later window when acute psychotomimetic effects had subsided but receptor-level modulation was likely ongoing (Duman et al. 2016; Zanos et al. 2016; Zanos et al. 2018; Kamp et al. 2020).

Our scanning time point (∼5 h post-infusion) corresponds to a subacute pharmacodynamic phase. This period occurs after the resolution of acute psychotomimetic symptoms (Ballard and Zarate 2020; Tashakkori et al. 2021) and coincides with the onset of antidepressant effects in MDD (Berman et al. 2000; Zarate et al. 2006). Mechanistically, this window is thought to involve glutamate-driven synaptic strengthening and circuit-level reorganisation, mediated by AMPA receptor activation, BDNF release, and mTOR signalling (Li et al. 2010; Duman et al. 2016; Zanos et al. 2016; Zanos et al. 2018; Duman et al. 2021).

Pharmacokinetic and pharmacodynamic data indicate that ketamine’s NMDA receptor antagonism peaks rapidly after administration and declines over several hours, consistent with a plasma elimination half-life of approximately 2 to 4 h (Mion and Villevieille 2013; Peltoniemi et al. 2016). Preclinical evidence suggests that ketamine may continue to block NMDA receptors for up to 24 h due to trapping mechanisms (Ma et al. 2023). Norketamine is eliminated more slowly and maintains high plasma concentrations more than 5 h after infusion (Quibell et al. 2011; Mion and Villevieille 2013). By the time of our scan (∼5 h post-infusion), NMDA receptor blockade has likely subsided (Zanos et al. 2018; Kamp et al. 2020), and secondary metabolites such as hydroxynorketamine predominate (Zanos et al. 2018). Although these metabolites are implicated in the sustained antidepressant effects of ketamine (Zanos et al. 2016), their immediate impact on task-evoked BOLD responses during reward processing remains unclear (Lally et al. 2014; Zanos et al. 2016).

The temporal mismatch between the peak of receptor-level effects and our task window may account for the absence of robust effects in our dataset. Ketamine’s effects on reward circuitry in healthy subjects may be time-specific, with fMRI alterations most reliably detected during the acute post-infusion window (<2 h) when NMDA receptor antagonism and dissociative effects are most pronounced (Lally et al. 2014; Francois et al. 2016; Kotoula et al. 2022). By approximately 5 h, when pharmacokinetics shift toward downstream metabolites and behavioral normalisation occurs, ketamine’s influence on reward processing may become more subtle, potentially manifesting through longer-term plasticity mechanisms rather than immediate modulation of task-related neural activity (Li et al. 2010; Duman et al. 2016; Zanos et al. 2016; Zanos et al. 2018; Duman et al. 2021). This intermediate state may therefore represent a transitional pharmacodynamic window between acute receptor-level effects and sustained synaptic changes. Future studies using denser temporal sampling across acute (≤2 h), intermediate (3–6 h), and later phases (24–72 h) could map evolving neural effects of ketamine more precisely.

It is also conceivable that the absence of ketamine-induced neural effects in our healthy cohort may result from baseline differences from MDD patients (Kotoula et al. 2022). MDD is associated with deficits in reward processing, including attenuated striatal and prefrontal responses (Zhang et al. 2013; Ng et al. 2019) and ketamine may differently modulate reward processing in individuals with pre-existing reward dysfunction. Indeed, ketamine-induced striatal activation has been reported in depressed individuals (Lally et al. 2014; Kotoula et al. 2022), though some studies have also reported ketamine-induced effects in healthy subjects (Francois et al. 2016; Mkrtchian et al. 2021). Mkrtchian et al. further demonstrated diagnosis-specific connectivity changes 48 h post-infusion, with ketamine normalising fronto-striatal connectivity in MDD but shifting it in healthy controls toward patterns typical of unmedicated MDD (Mkrtchian et al. 2021). These divergent effects suggest that ketamine may restore dysfunctional circuits in depression but disrupt the same networks in healthy individuals, consistent with state-dependent modulation (Reed et al. 2019; Mkrtchian et al. 2021). Evidence indicates that ketamine transiently induces symptoms related to impaired motivation in healthy participants (Driesen et al. 2013; Pollak et al. 2015), further implicating that the direction of ketamine’s effects on reward circuitry may be state-dependent.

### Limitations

Several limitations should be considered when interpreting these findings. First, our study included healthy subjects only. The lack of a substance effect on contrasts tested may not translate to clinical populations with pre-existing reward dysfunction, such as MDD patients. Second, the timing of the fMRI task (∼5 h post-infusion) could have missed the window in which ketamine exerts its most pronounced effects on reward processing. Although this time point was chosen to investigate subacute changes relevant to sustained antidepressant mechanisms, it may have fallen into a transitional pharmacodynamic state in which neither acute pharmacological effects nor long-term synaptic changes were sufficiently pronounced to produce detectable BOLD signal differences. Denser temporal sampling across acute, intermediate, and later phases would be necessary to fully capture the evolving neural effects of ketamine. Third, while we modelled individual ketamine exposure using AUC values up to task onset, we did not directly assess plasma concentrations of downstream metabolites such as (2R,6R)-hydroxynorketamine at the time of scanning. AUCs may further not sufficiently characterise the complex pharmacokinetics and -dynamics, especially, to model individual (perceived) response.

Finally, limitations in effect size sensitivity should be acknowledged. Whole-brain models estimated 322 to 366 resolution elements for the calculated contrasts, which are equivalent to adjusted type-I error probabilities ranging from 1.6e-4 to 1.4e-4 on voxel level. With 28 participants, six covariates, a power of 0.80, and a reliability of r = 0.21 (Demidenko et al. 2024 July 9) this would require detectable effect sizes of around f = 0.69 / d = 1.38 under repeated-measures ANOVA assumptions. Under the same assumptions, testing a single ROI would require a detectable effect size of f = 0.37 / d = 0.74. Since many different models and contrasts were calculated in the current work, these figures should be interpreted as rough estimates. Effect size calculations were performed in G*Power 3.1.9.7 (ANOVA: Repeated measures, within factors). While these estimates made it unlikely that effects would be detected at the voxel level, the usually more liberal cluster-level or ROI analyses might have detected sufficiently strong neural effects of ketamine.

## Conclusion

We observed no significant drug-related effects on reward processing during the MID task in healthy participants five hours after ketamine infusion. These findings combined with previous literature suggest that ketamine’s effects on reward circuitry are time-sensitive and potentially most prominent during the acute post-infusion phase (<2 h) (Lally et al. 2014; Francois et al. 2016; Kotoula et al. 2022). The 5 h post-infusion time point may reflect a transitional pharmacodynamic window, in which receptor-level effects have diminished but sustained synaptic changes have not yet fully developed. In this state, ketamine’s influence on anticipation- and feedback-related activation, as well as the Expectation–Outcome Congruency contrast, may be too subtle to detect. Moreover, ketamine’s neural effects may be state-dependent, with more robust modulation emerging in individuals with pre-existing reward system dysfunction, such as MDD patients (Lally et al. 2014; Kotoula et al. 2022; Nogo et al. 2022). Future studies should integrate longitudinal designs with denser temporal sampling across acute, intermediate, and later phases, and explicitly compare healthy and clinical cohorts to further characterise temporal dynamics and state-dependence of ketamine’s effects on reward processing.

## Acknowledgements

The authors would like to thank all staff involved in successful study implementation, including medical doctors and technical colleagues, and diploma students of the Advanced Neuroimaging Labs (aNIL, head: R. Lanzenberger), Department for Psychiatry and Psychotherapy (chair: D. Rujescu). They also express their gratitude to their collaboration partners at the Vienna Science Hub of the University of Vienna.

## Consent Declaration

All participants signed an informed consent at the time of enrolment and agreed to participate in the study.

## Data Availability Statement

Datasets of participants presented in this article are not readily available due to ethical reasons and data sharing regulations. Please contact for questions.

## References

Admon R, Pizzagalli DA. 2015. Dysfunctional Reward Processing in Depression. Curr Opin Psychol. 4:114–118. doi:10.1016/j.copsyc.2014.12.011.

Andersson JLR, Skare S, Ashburner J. 2003. How to correct susceptibility distortions in spin-echo echo-planar images: application to diffusion tensor imaging. Neuroimage. 20(2):870–888. doi:10.1016/S1053-8119(03)00336-7.

Argyropoulos SV, Nutt DJ. 2013. Anhedonia revisited: is there a role for dopamine-targeting drugs for depression? J Psychopharmacol. 27(10):869–877. doi:10.1177/0269881113494104.

Ballard ED, Zarate CA. 2020. The role of dissociation in ketamine’s antidepressant effects. Nat Commun. 11(1):6431. doi:10.1038/s41467-020-20190-4.

Balleine BW, O’Doherty JP. 2010. Human and Rodent Homologies in Action Control: Corticostriatal Determinants of Goal-Directed and Habitual Action. Neuropsychopharmacol. 35(1):48–69. doi:10.1038/npp.2009.131.

Beall EB, Lowe MJ. 2014. SimPACE: generating simulated motion corrupted BOLD data with synthetic-navigated acquisition for the development and evaluation of SLOMOCO: a new, highly effective slicewise motion correction. Neuroimage. 101:21–34. doi:10.1016/j.neuroimage.2014.06.038.

Behzadi Y, Restom K, Liau J, Liu TT. 2007. A component based noise correction method (CompCor) for BOLD and perfusion based fMRI. Neuroimage. 37(1):90–101. doi:10.1016/j.neuroimage.2007.04.042.

Berman RM, Cappiello A, Anand A, Oren DA, Heninger GR, Charney DS, Krystal JH. 2000. Antidepressant effects of ketamine in depressed patients. Biol Psychiatry. 47(4):351–354. doi:10.1016/s0006-3223(99)00230-9.

Bioanalytical method validation - Scientific guideline | European Medicines Agency (EMA). 2011 Aug 1. [accessed 2025 June 18]. https://www.ema.europa.eu/en/bioanalytical-method-validation-scientific-guideline.

Borsini A, Wallis ASJ, Zunszain P, Pariante CM, Kempton MJ. 2020. Characterizing anhedonia: A systematic review of neuroimaging across the subtypes of reward processing deficits in depression. Cogn Affect Behav Neurosci. 20(4):816–841. doi:10.3758/s13415-020-00804-6.

Carter RM, MacInnes JJ, Huettel SA, Adcock RA. 2009. Activation in the VTA and Nucleus Accumbens Increases in Anticipation of Both Gains and Losses. Front Behav Neurosci. 3:21. doi:10.3389/neuro.08.021.2009.

Casarotto PC, Girych M, Fred SM, Kovaleva V, Moliner R, Enkavi G, Biojone C, Cannarozzo C, Sahu MP, Kaurinkoski K, et al. 2021. Antidepressant drugs act by directly binding to TRKB neurotrophin receptors. Cell. 184(5):1299–1313.e19. doi:10.1016/j.cell.2021.01.034.

Chen Y, Chaudhary S, Li C-SR. 2022. Shared and distinct neural activity during anticipation and outcome of win and loss: A meta-analysis of the monetary incentive delay task. NeuroImage. 264:119764. doi:10.1016/j.neuroimage.2022.119764.

Costumero V, Barrós-Loscertales A, Bustamante JC, Ventura-Campos N, Fuentes P, Ávila C. 2013. Reward sensitivity modulates connectivity among reward brain areas during processing of anticipatory reward cues. European Journal of Neuroscience. 38(3):2399–2407. doi:10.1111/ejn.12234.

Coyle CM, Laws KR. 2015. The use of ketamine as an antidepressant: a systematic review and meta-analysis. Hum Psychopharmacol. 30(3):152–163. doi:10.1002/hup.2475.

Demidenko MI, Mumford JA, Poldrack RA. 2024 July 9. Impact of analytic decisions on test-retest reliability of individual and group estimates in functional magnetic resonance imaging: a multiverse analysis using the monetary incentive delay task. bioRxiv.:2024.03.19.585755. doi:10.1101/2024.03.19.585755.

Dittrich A. 1998. The standardized psychometric assessment of altered states of consciousness (ASCs) in humans. Pharmacopsychiatry. 31 Suppl 2:80–84. doi:10.1055/s-2007-979351.

Domino EF, Chodoff P, Corssen G. 1965. Pharmacologic effects of CI-581, a new dissociative anesthetic, in man. Clinical Pharmacology & Therapeutics. 6(3):279–291. doi:10.1002/cpt196563279.

Driesen NR, McCarthy G, Bhagwagar Z, Bloch M, Calhoun V, D’Souza DC, Gueorguieva R, He G, Ramachandran R, Suckow RF, et al. 2013. Relationship of resting brain hyperconnectivity and schizophrenia-like symptoms produced by the NMDA receptor antagonist ketamine in humans. Mol Psychiatry. 18(11):1199– 1204. doi:10.1038/mp.2012.194.

Dugré JR, Dumais A, Bitar N, Potvin S. 2018. Loss anticipation and outcome during the Monetary Incentive Delay Task: a neuroimaging systematic review and meta-analysis. PeerJ. 6:e4749. doi:10.7717/peerj.4749.

Duman RS, Aghajanian GK, Sanacora G, Krystal JH. 2016. Synaptic plasticity and depression: New insights from stress and rapid-acting antidepressants. Nat Med. 22(3):238–249. doi:10.1038/nm.4050.

Duman RS, Deyama S, Fogaça MV. 2021. Role of BDNF in the pathophysiology and treatment of depression: Activity-dependent effects distinguish rapid-acting antidepressants. Eur J Neurosci. 53(1):126–139. doi:10.1111/ejn.14630.

Ebert B, Mikkelsen S, Thorkildsen C, Borgbjerg FM. 1997. Norketamine, the main metabolite of ketamine, is a non-competitive NMDA receptor antagonist in the rat cortex and spinal cord. Eur J Pharmacol. 333(1):99–104. doi:10.1016/s0014-2999(97)01116-3.

Francois J, Grimm O, Schwarz AJ, Schweiger J, Haller L, Risterucci C, Böhringer A, Zang Z, Tost H, Gilmour G, et al. 2016. Ketamine Suppresses the Ventral Striatal Response to Reward Anticipation: A Cross-Species Translational Neuroimaging Study. Neuropsychopharmacol. 41(5):1386–1394. doi:10.1038/npp.2015.291.

Gorwood P. 2008. Neurobiological mechanisms of anhedonia. Dialogues Clin Neurosci. 10(3):291–299. doi:10.31887/DCNS.2008.10.3/pgorwood.

Hahn A, Reed MB, Murgaš M, Vraka C, Klug S, Schmidt C, Godbersen GM, Eggerstorfer B, Gomola D, Silberbauer LR, et al. 2025. Dynamics of human serotonin synthesis differentially link to reward anticipation and feedback. Mol Psychiatry. 30(2):600–607. doi:10.1038/s41380-024-02696-1.

Hahn A, Reed MB, Pichler V, Michenthaler P, Rischka L, Godbersen GM, Wadsak W, Hacker M, Lanzenberger R. 2021. Functional dynamics of dopamine synthesis during monetary reward and punishment processing. J Cereb Blood Flow Metab. 41(11):2973–2985. doi:10.1177/0271678X211019827.

Hahn T, Dresler T, Ehlis A-C, Plichta MM, Heinzel S, Polak T, Lesch K-P, Breuer F, Jakob PM, Fallgatter AJ. 2009. Neural response to reward anticipation is modulated by Gray’s impulsivity. Neuroimage. 46(4):1148–1153. doi:10.1016/j.neuroimage.2009.03.038.

Höflich A, Michenthaler P, Kasper S, Lanzenberger R. 2018. Circuit Mechanisms of Reward, Anhedonia, and Depression. Int J Neuropsychopharmacol. 22(2):105–118. doi:10.1093/ijnp/pyy081.

Jauhar S, Fortea L, Solanes A, Albajes-Eizagirre A, McKenna PJ, Radua J. 2021. Brain activations associated with anticipation and delivery of monetary reward: A systematic review and meta-analysis of fMRI studies. PLOS ONE. 16(8):e0255292. doi:10.1371/journal.pone.0255292.

Kamp J, Jonkman K, van Velzen M, Aarts L, Niesters M, Dahan A, Olofsen E. 2020. Pharmacokinetics of ketamine and its major metabolites norketamine, hydroxynorketamine, and dehydronorketamine: a model-based analysis. Br J Anaesth. 125(5):750–761. doi:10.1016/j.bja.2020.06.067.

Knutson B, Adams CM, Fong GW, Hommer D. 2001. Anticipation of increasing monetary reward selectively recruits nucleus accumbens. J Neurosci. 21(16):RC159. doi:10.1523/JNEUROSCI.21-16-j0002.2001.

Knutson B, Greer SM. 2008. Anticipatory affect: neural correlates and consequences for choice. Philosophical Transactions of the Royal Society B: Biological Sciences. 363(1511):3771–3786. doi:10.1098/rstb.2008.0155.

Knutson B, Westdorp A, Kaiser E, Hommer D. 2000. FMRI Visualization of Brain Activity during a Monetary Incentive Delay Task. NeuroImage. 12(1):20–27. doi:10.1006/nimg.2000.0593.

Kotoula V, Stringaris A, Mackes N, Mazibuko N, Hawkins PCT, Furey M, Curran HV, Mehta MA. 2022. Ketamine Modulates the Neural Correlates of Reward Processing in Unmedicated Patients in Remission From Depression. Biological Psychiatry: Cognitive Neuroscience and Neuroimaging. 7(3):285–292. doi:10.1016/j.bpsc.2021.05.009.

Kwaśny A, Cubała WJ, Włodarczyk A. 2024. Anhedonia and depression severity measures during ketamine administration in treatment-resistant depression. Front Psychiatry. 15:1334293. doi:10.3389/fpsyt.2024.1334293.

Kwaśny A, Kwaśna J, Wilkowska A, Szarmach J, Słupski J, Włodarczyk A, Cubała WJ. 2024. Ketamine treatment for anhedonia in unipolar and bipolar depression: a systematic review. European Neuropsychopharmacology. 86:20–34. doi:10.1016/j.euroneuro.2024.04.014.

Lally N, Nugent AC, Luckenbaugh DA, Ameli R, Roiser JP, Zarate CA. 2014. Anti-anhedonic effect of ketamine and its neural correlates in treatment-resistant bipolar depression. Translational Psychiatry. 4(10):e469. doi:10.1038/tp.2014.105.

Lally N, Nugent AC, Luckenbaugh DA, Niciu MJ, Roiser JP, Zarate CA. 2015. Neural correlates of change in major depressive disorder anhedonia following open-label ketamine. J Psychopharmacol. 29(5):596–607. doi:10.1177/0269881114568041.

Li N, Lee B, Liu R-J, Banasr M, Dwyer JM, Iwata M, Li X-Y, Aghajanian G, Duman RS. 2010. mTOR-Dependent Synapse Formation Underlies the Rapid Antidepressant Effects of NMDA Antagonists. Science. 329(5994):959–964. doi:10.1126/science.1190287.

Lumsden EW, Troppoli TA, Myers SJ, Zanos P, Aracava Y, Kehr J, Lovett J, Kim S, Wang F-H, Schmidt S, et al. 2019. Antidepressant-relevant concentrations of the ketamine metabolite (2R,6R)-hydroxynorketamine do not block NMDA receptor function. Proceedings of the National Academy of Sciences. 116(11):5160–5169. doi:10.1073/pnas.1816071116.

Lutz K, Widmer M. 2014. What can the monetary incentive delay task tell us about the neural processing of reward and punishment? NAN. 3:33–45. doi:10.2147/NAN.S38864.

Ma S, Chen M, Jiang Y, Xiang X, Wang S, Wu Z, Li S, Cui Y, Wang J, Zhu Y, et al. 2023. Sustained antidepressant effect of ketamine through NMDAR trapping in the LHb. Nature. 622(7984):802–809. doi:10.1038/s41586-023-06624-1.

Menon V, Uddin LQ. 2010. Saliency, switching, attention and control: a network model of insula function. Brain Struct Funct. 214(5–6):655–667. doi:10.1007/s00429-010-0262-0.

Mion G, Villevieille T. 2013. Ketamine pharmacology: an update (pharmacodynamics and molecular aspects, recent findings). CNS Neurosci Ther. 19(6):370–380. doi:10.1111/cns.12099.

Mkrtchian A, Evans JW, Kraus C, Yuan P, Kadriu B, Nugent AC, Roiser JP, Zarate CA. 2021. Ketamine modulates fronto-striatal circuitry in depressed and healthy individuals. Mol Psychiatry. 26(7):3292–3301. doi:10.1038/s41380-020-00878-1.

Murrough JW, Iosifescu DV, Chang LC, Al Jurdi RK, Green CE, Perez AM, Iqbal S, Pillemer S, Foulkes A, Shah A, et al. 2013. Antidepressant efficacy of ketamine in treatment-resistant major depression: a two-site randomized controlled trial. Am J Psychiatry. 170(10):1134–1142. doi:10.1176/appi.ajp.2013.13030392.

Ng TH, Alloy LB, Smith DV. 2019. Meta-analysis of reward processing in major depressive disorder reveals distinct abnormalities within the reward circuit. Transl Psychiatry. 9(1):293. doi:10.1038/s41398-019-0644-x.

Nikolin S, Rodgers A, Schwaab A, Bahji A, Zarate C, Vazquez G, Loo C. 2023. Ketamine for the treatment of major depression: a systematic review and meta-analysis. EClinicalMedicine. 62:102127. doi:10.1016/j.eclinm.2023.102127.

Nogo D, Jasrai AK, Kim H, Nasri F, Ceban F, Lui LMW, Rosenblat JD, Vinberg M, Ho R, McIntyre RS. 2022. The effect of ketamine on anhedonia: improvements in dimensions of anticipatory, consummatory, and motivation-related reward deficits. Psychopharmacology (Berl). 239(7):2011–2039. doi:10.1007/s00213-022-06105-9.

Nugent AC, Diazgranados N, Carlson PJ, Ibrahim L, Luckenbaugh DA, Brutsche N, Herscovitch P, Drevets WC, Zarate CA. 2014. Neural correlates of rapid antidepressant response to ketamine in bipolar disorder. Bipolar Disord. 16(2):119–128. doi:10.1111/bdi.12118.

Oldham S, Murawski C, Fornito A, Youssef G, Yücel M, Lorenzetti V. 2018. The anticipation and outcome phases of reward and loss processing: A neuroimaging meta-analysis of the monetary incentive delay task. Human Brain Mapping. 39(8):3398–3418. doi:10.1002/hbm.24184.

Patel AX, Bullmore ET. 2016. A wavelet-based estimator of the degrees of freedom in denoised fMRI time series for probabilistic testing of functional connectivity and brain graphs. NeuroImage. 142:14–26. doi:10.1016/j.neuroimage.2015.04.052.

Patel AX, Kundu P, Rubinov M, Jones PS, Vértes PE, Ersche KD, Suckling J, Bullmore ET. 2014. A wavelet method for modeling and despiking motion artifacts from resting-state fMRI time series. Neuroimage. 95(100):287–304. doi:10.1016/j.neuroimage.2014.03.012.

Peltoniemi MA, Hagelberg NM, Olkkola KT, Saari TI. 2016. Ketamine: A Review of Clinical Pharmacokinetics and Pharmacodynamics in Anesthesia and Pain Therapy. Clin Pharmacokinet. 55(9):1059–1077. doi:10.1007/s40262-016-0383-6.

Pizzagalli DA, Holmes AJ, Dillon DG, Goetz EL, Birk JL, Bogdan R, Dougherty DD, Iosifescu DV, Rauch SL, Fava M. 2009. Reduced caudate and nucleus accumbens response to rewards in unmedicated individuals with major depressive disorder. Am J Psychiatry. 166(6):702–710. doi:10.1176/appi.ajp.2008.08081201.

Pollak TA, De Simoni S, Barimani B, Zelaya FO, Stone JM, Mehta MA. 2015. Phenomenologically distinct psychotomimetic effects of ketamine are associated with cerebral blood flow changes in functionally relevant cerebral foci: a continuous arterial spin labelling study. Psychopharmacology (Berl). 232(24):4515–4524. doi:10.1007/s00213-015-4078-8.

Pulcu E, Guinea C, Cowen PJ, Murphy SE, Harmer CJ. 2022. A translational perspective on the anti-anhedonic effect of ketamine and its neural underpinnings. Mol Psychiatry. 27(1):81–87. doi:10.1038/s41380-021-01183-1.

Quibell R, Prommer EE, Mihalyo M, Twycross R, Wilcock A. 2011. Ketamine*. Journal of Pain and Symptom Management. 41(3):640–649. doi:10.1016/j.jpainsymman.2011.01.001.

Reed JL, Nugent AC, Furey ML, Szczepanik JE, Evans JW, Zarate CA. 2019. Effects of Ketamine on Brain Activity During Emotional Processing: Differential Findings in Depressed Versus Healthy Control Participants. Biological Psychiatry: Cognitive Neuroscience and Neuroimaging. 4(7):610–618. doi:10.1016/j.bpsc.2019.01.005.

Rolls ET, Huang C-C, Lin C-P, Feng J, Joliot M. 2020. Automated anatomical labelling atlas 3. NeuroImage. 206:116189. doi:10.1016/j.neuroimage.2019.116189.

Sałat K, Siwek A, Starowicz G, Librowski T, Nowak G, Drabik U, Gajdosz R, Popik P. 2015. Antidepressant-like effects of ketamine, norketamine and dehydronorketamine in forced swim test: Role of activity at NMDA receptor. Neuropharmacology. 99:301–307. doi:10.1016/j.neuropharm.2015.07.037.

Schultz W. 2016. Dopamine reward prediction-error signalling: a two-component response. Nat Rev Neurosci. 17(3):183–195. doi:10.1038/nrn.2015.26.

Schultz W, Dayan P, Montague PR. 1997. A neural substrate of prediction and reward. Science. 275(5306):1593–1599. doi:10.1126/science.275.5306.1593.

Seeley WW, Menon V, Schatzberg AF, Keller J, Glover GH, Kenna H, Reiss AL, Greicius MD. 2007. Dissociable Intrinsic Connectivity Networks for Salience Processing and Executive Control. J Neurosci. 27(9):2349–2356. doi:10.1523/JNEUROSCI.5587-06.2007.

Shin W, Koenig KA, Lowe MJ. 2022. A comprehensive investigation of physiologic noise modeling in resting state fMRI; time shifted cardiac noise in EPI and its removal without external physiologic signal measures. Neuroimage. 254:119136. doi:10.1016/j.neuroimage.2022.119136.

Smoski MJ, Felder J, Bizzell J, Green SR, Ernst M, Lynch TR, Dichter GS. 2009. fMRI of alterations in reward selection, anticipation, and feedback in major depressive disorder. J Affect Disord. 118(1–3):69–78. doi:10.1016/j.jad.2009.01.034.

Spijker J, Bijl RV, de Graaf R, Nolen WA. 2001. Determinants of poor 1-year outcome of DSM-III-R major depression in the general population: results of the Netherlands Mental Health Survey and Incidence Study (NEMESIS). Acta Psychiatr Scand. 103(2):122–130. doi:10.1034/j.1600-0447.2001.103002122.x.

Studerus E, Gamma A, Vollenweider FX. 2010. Psychometric Evaluation of the Altered States of Consciousness Rating Scale (OAV). PLoS One. 5(8):e12412. doi:10.1371/journal.pone.0012412.

Szczepanik JE, Reed JL, Nugent AC, Ballard ED, Evans JW, Lejuez CW, Zarate CA. 2019. Mapping anticipatory anhedonia: an fMRI study. Brain Imaging Behav. 13(6):1624–1634. doi:10.1007/s11682-019-00084-w.

Tashakkori M, Ford A, Dragovic M, Gabriel L, Waters F. 2021. The time course of psychotic symptom side effects of ketamine in the treatment of depressive disorders: a systematic review and meta-analysis. Australas Psychiatry. 29(1):80–87. doi:10.1177/1039856220961642.

Wilson RP, Colizzi M, Bossong MG, Allen P, Kempton M, Abe N, Barros-Loscertales AR, Bayer J, Beck A, Bjork J, et al. 2018. The Neural Substrate of Reward Anticipation in Health: A Meta-Analysis of fMRI Findings in the Monetary Incentive Delay Task. Neuropsychol Rev. 28(4):496–506. doi:10.1007/s11065-018-9385-5.

Winer ES, Nadorff MR, Ellis TE, Allen JG, Herrera S, Salem T. 2014. Anhedonia predicts suicidal ideation in a large psychiatric inpatient sample. Psychiatry Res. 218(1–2):124–128. doi:10.1016/j.psychres.2014.04.016.

Zanos P, Moaddel R, Morris PJ, Georgiou P, Fischell J, Elmer GI, Alkondon M, Yuan P, Pribut HJ, Singh NS, et al. 2016. NMDAR inhibition-independent antidepressant actions of ketamine metabolites. Nature. 533(7604):481–486. doi:10.1038/nature17998.

Zanos P, Thompson SM, Duman RS, Zarate CA, Gould TD. 2018. Convergent Mechanisms Underlying Rapid Antidepressant Action. CNS Drugs. 32(3):197–227. doi:10.1007/s40263-018-0492-x.

Zarate CA, Singh JB, Carlson PJ, Brutsche NE, Ameli R, Luckenbaugh DA, Charney DS, Manji HK. 2006. A randomized trial of an N-methyl-D-aspartate antagonist in treatment-resistant major depression. Arch Gen Psychiatry. 63(8):856–864. doi:10.1001/archpsyc.63.8.856.

Zhang W-N, Chang S-H, Guo L-Y, Zhang K-L, Wang J. 2013. The neural correlates of reward-related processing in major depressive disorder: a meta-analysis of functional magnetic resonance imaging studies. J Affect Disord. 151(2):531–539. doi:10.1016/j.jad.2013.06.039.

